# Tomato SlBES1.8 influences leaf morphogenesis by mediating gibberellin metabolism and signaling

**DOI:** 10.1101/2021.12.10.472069

**Authors:** Deding Su, Wei Xiang, Qin Liang, Ling Wen, Yuan Shi, Yudong Liu, Zhiqiang Xian, Zhengguo Li

## Abstract

Leaf morphogenetic activity determines its shape diversity. However, our knowledge to the regulatory mechanism in maintaining leaf morphogenetic capacity is still limited. In tomato, gibberellin (GA) negatively regulates leaf complexity by shortening the morphogenetic window. We here reported a tomato BRI1-EMS-SUPPRESSOR 1 (BES1) transcription factor, SlBES1.8, that promoted the simplification of leaf pattern in a similar manner as GA functions. Enhanced level of SlBES1.8 dramatically decreased the sensibility of tomato to GA whereas increased the sensibility to the GA biosynthesis inhibitor, PAC. In line with the phenotypic observation, the endogenous bioactive GA contents were increased in OE-*SlBES1*.*8* lines, which certainly promoted the degradation of the GA signaling negative regulator, SlDELLA. Moreover, transcriptomic analysis uncovered a set of overlapping genomic targets of SlBES1.8 and GA, and most of them were regulated in the same way. Expression studies showed the repression of SlBES1.8 to the transcriptions of two GA deactivated genes, *SlGA2ox2* and *SlGA2ox6*, and one GA receptor, *SlGID1b-1*. Further experiments confirmed the direct regulation of SlBES1.8 to their promoters. On the other hand, SlDELLA physically interacted with SlBES1.8 and further inhibited its transcriptional regulation activity by abolishing SlBES1.8-DNA binding. Conclusively, by mediating GA deactivation and signaling, SlBES1.8 greatly influenced tomato leaf morphogenesis.

**Highlight:** The BRI1-EMS-SUPPRESSOR 1 (BES1) family protein SlBES1.8 promotes leaf simplification by inhibiting gibberellin deactivation and physically interacting with DELLA protein in tomato.

## Introduction

Leaf is a vital plant organ not only because of its functions as a solar panel that utilizing light energy to converts carbon dioxide into carbohydrates, but also for the integration of internal and external signals in response to various environmental information (Du *et al*., 2018). Leaf primordia initiate from the peripheral zone (PZ) of shoot apical meristem (SAM), then the three axes of polar growth, that are the adaxial-abaxial, proximal-distal and medio-lateral axes, are gradually established (Byrne, 2012; Du *et al*., 2018). The margins of leaf primordia maintain a transient window of meristematic activity, allowing the production of leaflets, lobes and serrations, thus fine-tuning of this window enables the variability in leaf patterning (Alvarez *et al*., 2016; Israeli *et al*., 2021). According to the number of blade, leaves are classified as simple leaf that only has a single blade and compound leaf that owns several separate blades attached to a common rachis (Efroni *et al*., 2010). Tomato (*Solanum lycopersicum*) is a typical model plant that possesses compound leaves, which are composed of four kinds of leaflets: terminal leaflet (TL), primary lateral leaflet (PLL), secondary lateral leaflet (SLL) and intercalary leaflet (IL), and the lobes or serrations can be found in the margin of leaflet. There are usually one TL and three pairs of PLLs in tomato leaf branch. SLL develops from the petiolule of PLL, whereas IL develops later from the rachis between PLL, and their numbers are diverse and indeterminate (Goliber *et al*., 1999).

The initiation, differentiation and maturation of plant leaf are controlled by the combination of transcriptional regulators, microRNAs and phytohormones. The initiation of leaf primordia is required for the presence of auxin maxima, which are established by the auxin efflux carrier PINFORMED1 (PIN1) (Reinhardt *et al*., 2000; Reinhardt *et al*., 2003). Late study reported that auxin induced the expression of a tomato ortholog of Arabidopsis *DORNRONSCHEN* (*DRN*) gene, *LEAFLESS* (*LFS*), which is necessary for leaf initiation, to specify primordium cells in the PZ (Capua and Eshed, 2017). Besides, many transcription factors are involved in the regulation of leaf initiation, including class I KNOTTED1-like homeobox proteins (KNOXI) and the ASYMMETRIC LEAVES1 (AS1)/ROUGH SHEATH2 (RS2)/PHANTASTICA (ARP) MYB-domain transcription factors (Kerstetter *et al*., 1997; Byrne *et al*., 2000; Lodha *et al*., 2013). During leaf patterning, a set of regulating factors including MONOPTEROS (MP, also named ARF5), GOBLET (GOB), LYRATE (LYR) and REDUCED COMPLEXITY (RCO) coordinately modulate the formation and shaping of leaflet in parallel different aspects (Vlad *et al*., 2014; Israeli *et al*., 2021). Leaf patterning is also greatly regulated by several miRNA-target modules. For instance, miR165/166 contributes to the establishment of leaf adaxial-abaxial polarity by targeting *HD-ZIP* □ mRNAs (Tatematsu *et al*., 2015; Merelo *et al*., 2016), miR319 and miR396 antagonistically regulate leaf marginal cell proliferation by targeting the class □ TEOSINTE BRANCHED1/CYCLOIDEA/PROLIFERATING CELL FACTOR (TCP) and GROWTH-REGULATING FACTORs (GRFs) respectively (Ori *et al*., 2007; Rodriguez *et al*., 2010; Tsukaya, 2018).

Leaf morphogenesis is also regulated by several phytohormones (Shwartz *et al*., 2016). Among them, Gibberellin (GA) is shown to reduce the complexity of tomato leaf. Exogenous GA application promotes the maturation of tomato leaf and inhibits the formation of leaflets. Similarly, leaves of the *procera* (*pro*) mutant, which sustained mutation in the single tomato DELLA gene and showed constitutive GA response, bear fewer leaflets with smooth margins (Jasinski *et al*., 2008; Lombardi-Crestana *et al*., 2012; Shwartz *et al*., 2016). This leaf simplification caused by increased GA levels or responses indicates the role of GA in shortening the morphogenetic window during leaf formation. Some leaf morphogenesis regulators are shown to function in mediating GA dynamics. In tomato, the CIN-TCP transcription factor LANCEOLATE (LA) accelerates leaf maturation and simplifies leaf shape partially by positively regulating GA levels (Yanai *et al*., 2011). By contrast, KNOXI protein antagonizes GA activity by repressing the expression of GA biosynthesis gene *SlGA20ox1* in regulating leaf morphogenesis (Jasinski *et al*., 2008).

In addition to GAs, Brassinosteroids (BRs) are another kind of plant growth-promoting hormone with functions in leaf development by promoting cell elongation and differentiation (Mitchell *et al*., 1970; Grove *et al*., 1979). BR biosynthetic and signaling mutants indeed show abnormal leaf patterning (Koka *et al*., 2000; Zhiponova *et al*., 2013). The crosstalk between GAs and BRs was well explored in the last decade. In Arabidopsis, the negative regulators of GA signaling, DELLA proteins, physically interacted with the dephosphorylated BRASSINAZOLE RESISTANT1 (BZR1), the key positive regulator of BR signaling (Bai *et al*., 2012; Gallego-Bartolomé *et al*., 2012; Li *et al*., 2012). Moreover, the transcriptional regulation activity of BZR1 was inhibited by the interaction with DELLA proteins (Bai *et al*., 2012; Gallego-Bartolomé *et al*., 2012; Li *et al*., 2012). On the other hand, genetic and biochemical evidences suggested that BZR1 and BRI1-EMS-SUPPRESSOR 1 (BES1) affected GA levels by directly regulating the expression of GA biosynthetic genes *GA20ox* and *GA3ox* in Arabidopsis and rice (Tong *et al*., 2014; Unterholzner *et al*., 2015). This integration of GA and BR both on signaling level and hormone level allowed the elaborated regulation of plant growth and leaf development.

Here, we reported a tomato BES1 family member, SlBES1.8, that can influence leaf morphogenesis by mediating GA deactivation and signaling. Overexpression of *SlBES1*.*8* in tomato significantly reduced the leaf complexity, which mimicked the functions of GA in regulating leaf development. Indeed, the endogenous bioactive GA contents were increased in OE-*SlBES1*.*8* leaves because of the transcriptional repression of *SlGA2ox2* and *SlGA2ox6* by SlBES1.8, which further resulted in the degradation of SlDELLA, hence the GA signaling was enhanced in OE-*SlBES1*.*8* lines. On the other hand, the transcriptional regulation ability of SlBES1.8 was inhibited by SlDELLA interaction, and such inhibition was released by the SlBES1.8-promoted degradation of SlDELLA, which further enhanced GA signaling in OE-*SlBES1*.*8* lines. Together, our results provide new insight into the crosstalk between BES1 transcription factors and GA levels or responses in regulating leaf morphogenesis.

## Materials and Methods

### Plant material growth conditions and generation of SlBES1.8 over-expressed lines

The coding sequence of *SlBES1*.*8* was cloned into K303 binary vector under the promotion of CaMV 35S promoter (Ren *et al*., 2011; Liu *et al*., 2020). The recombined expression vector was transformed into *Agrobacterium tumefaciens* strain GV3101 followed by cotyledon transformation according to V J Chetty *et al*. (2013) in tomato (*Solanum lycopersicum*) cv. Micro-Tom and *Ailsa Craig* background to generate the OE-*SlBES1*.*8* lines. Positive transgenic lines were screened by kanamycin and PCR detection, and the relative mRNA level of *SlBES1*.*8* in positive lines was detected by quantitative real-time polymerase chain reaction (qRT-PCR) in T2 generation. All tomato plants were cultivated in greenhouse with 16/8 h light/dark cycle, 25/20 □ day/night temperature and 60% relative humidity.

### Observation of leaf pattern by scanning electron microscopy (SEM)

The extra leaves of WT and OE-*SlBES1*.*8* plants at 15 DPG were hand-dissected to expose the leaf primordium. The shoots of seedlings were subsequently immersed into FAGE solution (50% absolute ethanol, 10% formaldehyde, 5% acetic acid and 0.72% glutaraldehyde) immediately for one day followed by dehydrating in an ethanol series from 30% to 100% over five days. The prepared samples were critically point dried and coated with gold. Finally, the images were taken by a FEI Inspect S50 scanning electron microscope.

### Gibberellin A3 (GA_3_) and paclobutrazol (PAC) treatment

GA_3_ and PAC (Sigma, USA) were dissolved in absolute methanol. For GA_3_ and PAC treatment, tomato seeds were germinated on MS/2 medium containing GA_3_ (10 μM and 50 μM) or PAC (5 μM) respectively after sterilization. The control seeds were germinated on MS/2 medium containing the same solution without GA_3_ and PAC. After 10-days normal growth, the seedlings were transplanted to the soil and sprayed with 100 μM GA_3_ or 50 μM PAC solution twice a week. Plant samples were observed or collected at the corresponding times as described in the context.

### Measurement of endogenous GAs contents

For measuring endogenous GAs contents, leaves of WT and OE-*SlBES1*.*8* plants at 5-week-old were harvested and frozen into liquid nitrogen immediately. The measurement was conducted by Wuhan Metware Biotechnology Co., Ltd. (Wuhan, China) based on the AB Sciex QTRAP 6500 LC-MS/MS platform. Briefly, leaf samples were ground using a mixer mill (MM 400, Retsch, Germany) to powder. GAs were extracted with 70% (v/v) acetonitrile from 500 mg powder. After adding the internal standards, 100 μL trimethylamine and 100 μL BPTAB, the resulting solution was vortexed, incubated at 90□ for 1 hour, and evaporated to dryness under nitrogen gas stream followed by redissolving in 1 mL 70% (v/v) acetonitrile. The extracts were filtered through 0.22 μm filter and then analyzed using an UPLC-ESI-MS/MS system (UPLC, ExionLCTM AD, https://sciex.com.cn/; MS, Applied Biosystems 6500 Triple Quadrupole, https://sciex.com.cn/). Finally, the output data was analyzed by Analyst 1.6.3 software.

### RNA extraction, cDNA synthesis and qRT-PCR analysis

Total RNA was extracted by using the RNeasy Plant Mini Kit (Qiagen, Germany) and treated with RNase-free DNase (Qiagen, Germany). The integrity and concentration of total RNA were detected by agarose gel electrophoresis and NanoDrop 1000 (Thermo, USA) respectively. After that, the PrimeScript™ RT reagent Kit with gDNA Eraser (TAKARA, Japan) was used to synthesize the first-strand cDNA with 1 μg total RNA as the template. The cDNA products were diluted to 5-fold with deionized water before use. qRT-PCR was performed by TB Green® Premix Ex Taq™ □ (Tli RNaseH Plus) (TAKARA, Japan) on the CFX96 Touch™ Real-Time PCR Detection System (BIO-RAD, USA) according to the manufacturer’s instructions. The relative fold change was calculated by the 2^-ΔΔCt^ method. *SlUBI* (Solyc01g056940) was used as the internal reference gene and all primers used in qRT-PCR detection were listed in Supplementary Table S1.

### RNA sequencing and data processing

The fourth leaf primordium (P4) of WT and OE-*SlBES1*.*8* seedlings at 16 DPG (that is L6 presented in Fig. 2) treated with or without 50 μM GA_3_ was microdissected with a surgical blade and frozen into liquid nitrogen immediately, which was regarded as the leaf samples used in RNA sequencing. At least 30 leaf primordia were collected for each sample, and three independent samples were sequenced for each group. The total RNA was extracted by using the PicoPure™ RNA Extraction kit (ARCTURUS, USA) according to the user guide. The concentration of total RNA was measured by NanoDrop 1000 (Thermo, USA). The integrity of total RNA was detected by agarose gel electrophoresis and 2100 Bioanalyser (Agilent, USA) and only those RNA samples showed clear bands and high RNA Integrity Number were conducted sequencing library. The RNA-seq was carried out in Majorbio (Shanghai Majorbio Bio-pharm Technology, Ltd, China) with the Illumina HiSeq xten/Novaseq 6000 sequencer. The raw data were trimmed and qualified by SeqPrep and Sickle with default parameters. The generated clean reads were subsequently mapped to the tomato genome in the Solanaceae Genomics Network (http://solgenomics.net/) by using HISAT2 software. Finally, DESeq2 was used to analyze the differentially expressed genes (DEGs) with fold change (FC) > 2 and adjust *P*-value (*Q*-value) < 0.05.

### Electrophoretic mobility shift assay (EMSA)

Full length of *SlBES1*.*8* and *SlDELLA* coding sequences were cloned into pGEX-4T-1 plasmid. The recombined plasmids were expressed in *Escherichia coli* strain Rosetta2 BL21 cells (Transgen, China) to produce the GST fused proteins. The fused proteins were purified with the GST-tagged protein purification kit (Clontech, USA) according to the manufacturer’s instructions. The designed probes of *SlGA2ox2, SlGA2ox6* and *SlGID1b-1* were biotin labeled following the instruction of Light Shift^®^ Chemiluminescent EMSA Kit (Thermo, USA). The mutated biotin probes that had an AAAAAA fragment instead of G-box were used to confirm the specific binding and the unlabeled probes were used as the competitor. For assessing the effect of SlDELLA to the DNA binding ability of SlBES1.8, SlBES1.8 and SlDELLA proteins were mixed before EMSA reaction. All native and mutated probes were listed in Supplementary Table S1.

### Yeast one-hybrid (Y1H) assays

The promoter fragments of *SlGA2ox2, SlGA2ox6* and *SlGID1b-1* were constructed into pAbAi plasmid and transformed into Y1HGold yeast strain under the guidance of Yeastmaker™ Yeast Transformation System 2 (TAKARA, Japan). The recombinant yeast strain was confirmed by PCR conducted with Matchmaker™ Insert Check PCR Mix (TAKARA, Japan). The full length of *SlBES1*.*8* coding sequence was cloned into pGADT7 plasmid and subsequently transformed into the recombinant yeast strain. The empty pGADT7 plasmid was used as negative control. The transformants were cultivated on SD/-Leu or SD/-Leu/AbA (Aureobasidin A, Clontech, USA) medium for three days. DNA-protein interactions between SlBES1.8 and promoter fragments were determined by the growth of colony.

### Dual-luciferase assays

To explore the direct transcriptional regulation of SlBES1.8 to *SlGA2ox2, SlGA2ox6* and *SlGID1b-1*, full length of *SlBES1*.*8* coding sequence was cloned into pGreen □ 62-SK to generate the effector, and promoter fragments of *SlGA2ox2, SlGA2ox6* and *SlGID1b-1* were constructed into pGreen II 0800-LUC respectively to generate the reporters (Hellens *et al*., 2005). After transiently co-expressed the effector and reporter in tobacco leaf for one-day darkness and another two-days normal growth, the expression of *LUC* and *REN* were detected by the Dual-Luciferase Reporter Assay System (Promega, USA). At least six biological replicates were measured, and the detection was performed independently for two times with similar results.

To explore the effect of SlDELLA on the transcriptional repression activity of SlBES1.8, full length of *SlBES1*.*8* and *SlDELLA* coding sequences were cloned into pEAQ-GAL4BD vector respectively to generate the effectors. In reporter, the expression of firefly luciferase (LUC) was under the control of GAL4-binding element (5×GAL4), which was fused with the minimal TATA region of CaMV35S, and the renilla luciferase (REN) promoted by CaMV35S was considered as the internal control. The SlBES1.8 and SlDELLA effectors were separately or together co-expressed with the reporter in tobacco leaves. Finally, the *LUC* and *REN* expression were detected by the Dual-Luciferase Reporter Assay System (Promega, USA). At least six biological replicates were measured, and the detection was performed independently for two times with similar results.

### Immunoblot assay

For immunoblot analyses, 16-days-old tomato seedlings were sprayed with 100 μM GA_3_ or 50 μM PAC solution for 3 hours, and then frozen into liquid nitrogen immediately. Protein extracts were performed with the Plant Total Protein Extraction Kit (Sigma, USA). Protein concentrations were measured using the BCA method. Equal amounts of protein from each sample were boiled in 4× Laemmli protein sample buffer (Bio-rad, USA) for 10 min followed by separating with 10% SDS-PAGE. After a wet transfer to PVDF membranes, the anti-GAI rabbit antibody and goat anti-rabbit IgG antibody (Agrisera, Sweden) were used to perform immunoblot.

### Yeast two-hybrid (Y2H) assays

Full length of *SlBES1*.*8* and *SlDELLA* coding sequences were constructed into pGBKT7 and pGADT7 as bait and prey respectively. The recombined plasmids were co-transformed into Y2H Gold yeast cells. Culture mediums SD/-Leu/-Trp and SD/-Ade/-His/-Leu/-Trp (Clontech, USA) with or without X-α-gal were used to select the positive transformants.

### Bimolecular fluorescence complementation (BiFC) assays

Full length of *SlBES1*.*8* and *SlDELLA* coding sequences were cloned into pXY104-YFP^C^ and pXY106-YFP^N^ (Yu *et al*., 2008) respectively. The BiFC assays were performed as described previously (Luo *et al*., 2014). In short, the recombined plasmids were transformed into *Agrobacterium tumefaciens* strain GV3101 and co-injected into and expressed in tobacco epidermic cells. The recombined plasmids co-injected with corresponding empty YFP^N^ or YFP^C^ were used as negative control. After incubated in darkness for one day and in normal light condition for another two days, the fluorescence images were captured by the laser scanning confocal microscope (Leica TCS SP8, Germany).

### Firefly luciferase complementation imaging (LCI) assays

Full length or truncated versions of *SlBES1*.*8* and *SlDELLA* were fused with nLUC or cLUC respectively. The LCI assays were conducted following the description by Chen *et al*. (2008). Briefly, the recombined plasmids were transformed into *Agrobacterium tumefaciens* strain GV3101. The fused SlBES1.8-nLUC transformants were transiently co-expressed with cLUC-SlDELLA transformants in tobacco leaves. After incubated in darkness for one day and in normal light condition for another two days, the tobacco leaves were sprayed with one millimolar luciferin (Promega, USA) on the blade back and kept in dark at least for six minutes. Finally, the LCI images were captured by a low-light cooled CCD imaging apparatus (Alliance, UK). At least four leaves were observed for each combination.

### Accession numbers

Arabidopsis genes are obtained from The Arabidopsis Information Resource (TAIR, https://www.arabidopsis.org/) under the following identities: *GAI*, AT1G14920; *RGA*, AT2G01570; *RGL1*, AT1G66350; *RGL2*, AT3G03450; *RGL3*, AT5G17490. Tomato genes are collected from Solanaceae Genomics Database (http://solgenomics.net/) under the following accession numbers: *SlBES1*.*8*, Solyc10g076390; *SlGA2ox2*, Solyc07g056670; *SlGA2ox6*, Solyc01g058030; *SlGID1b-1*, Solyc09g074270; *SlDELLA*, Solyc11g011260; *SlGA2ox1*, Solyc05g053340; *SlGA2ox3*, Solyc01g079200; *SlGA2ox4*, Solyc07g061720; *SlGA2ox5*, Solyc07g061730; *SlGA2ox7*, Solyc02g070430; *SlGA2ox8*, Solyc08g016660; *SlGA2ox9*, Solyc10g007570; *SlGID1ac*, Solyc01g098390; *SlGID1b-2*, Solyc06g008870; *SlSLY1*, Solyc04g078390; *SlSNE*, Solyc07g047680; *SlUBI*, Solyc01g056940. The raw RNA-sequencing data are publicly available in the Genome Sequence Archive in National Genomics Data Center, Beijing Institute of Genomics, Chinese Academy of Sciences with the accession number of CRA005324.

## Results

### SlBES1.8 mimics the effect of exogenous GA application in simplifying leaf complexity

Previously, we reported that *SlBES1*.*8* exhibited distinct gene structure compared to other *BES1* members in tomato and acted as a transcriptional repressor with floral tissues specific expression pattern (Su *et al*., 2021). To explore the functions of SlBES1.8 in plant development, we generated over-expressed (OE) lines driving by CaMV35S promoter in micro-tom background. Three OE-*SlBES1*.*8* lines (L4, L5 and L8) with dramatically increased *SlBES1*.*8* expressions were selected to analyze the phenotypic changes (Supplementary Fig. S1A). Remarkably, OE-*SlBES1*.*8* plants exhibited simpler leaf shape with smooth margin compared with wild type (WT) plants, and this phenotypic change occurred in early leaf primordium development (Fig. 1A-C). In WT leaf primordium, serrations were emerged from the margin of leaflet, while there was no such structure in OE-*SlBES1*.*8* leaf primordium (Fig. 1A, B). Consistent with this, the serration numbers of mature terminal leaflets and primary lateral leaflets were significantly decreased in OE-*SlBES1*.*8* leaves (Fig. 1C). Given that micro-tom is a dwarf cultivar of tomato and its leaves are simplified by the mutation in the *SELF-PRUNING* (*SP*) and *DWARF* (*D*) genes, which are related with shoot apical meristem development and BR biosynthesis respectively (Martí *et al*., 2006), we also generated OE-*SlBES1*.*8* lines in *Ailsa Craig* background (Supplementary Fig. S1B), in which the leaf pattern remained complicated. Consistent with the observation in micro-tom background, OE-*SlBES1*.*8* leaves also showed simpler shape and smooth margin (Fig. 1D, Supplementary Fig. S2). The leaflets can be briefly classified into three groups: terminal leaflet and primary lateral leaflets (1st), secondary lateral leaflets (2nd) and intercalary leaflets (3rd) (Fig. 1E). The numbers of all of these three kinds of leaflets were decreased in OE-*SlBES1*.*8* leaves (Fig. 1F). On the contrary, we also generated *slbes1*.*8* mutants by CRISPR/Cas9 technology. Two homozygous lines with distinct mutations were selected to analyze the leaf pattern (Supplementary Fig. S3A), while no obvious phenotypic changes were observed (Supplementary Fig. S3B), and the leaflet serration numbers of the 5^th^ leaf were not significantly influenced in *slbes1*.*8* mutants (Supplementary Fig. S3C), indicating that knockout of *SlBES1*.*8* didn’t affect the normal growth of tomato leaf development.

**Fig. 1.**
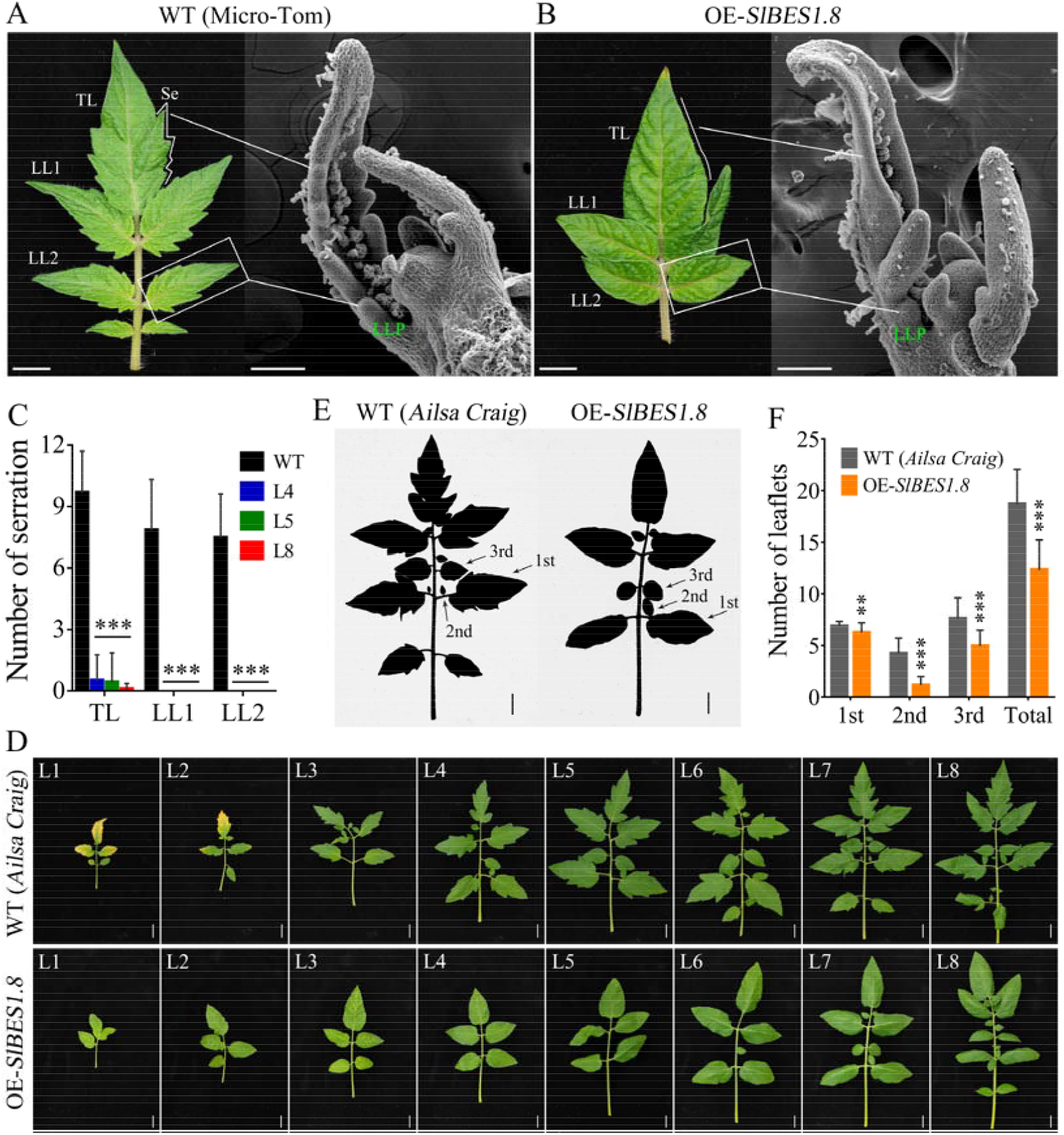
Overexpression of *SlBES1*.*8* resulted in changed morphogenesis of tomato leaf. (A, B) The mature leaf at 35 DPG and leaf primordium at 15 DPG of WT (A) and OE-*SlBES1*.*8* (B) plants in micro-tom background. TL, terminal leaflet; LL, lateral leaflet; LLP, lateral leaflet primordium; Se, serration; DPG, days post germination; bar in mature leaf indicates 1 cm, bar in leaf primordium indicates 100 μm. (C) Serration numbers of terminal leaflet and lateral leaflet in mature leaves. At least 20 leaves were counted for each column. (D) Leaves of wild type and OE-*SlBES1*.*8* plants in *Ailsa Craig* background. L, leaf; bar = 2 cm. (E) Silhouettes of the seventh leaf from wild type and OE-*SlBES1*.*8* plants (*Ailsa Craig* background) at 45 DPG. 1st indicates terminal and primary lateral leaflets; 2nd indicates secondary lateral leaflets; 3rd indicates intercalary leaflets; bar = 2 cm. (F) Leaflet numbers of wild type and OE-*SlBES1*.*8* plants presented in (D). At least 20 leaves were counted for each column. ** and *** refer to significant differences between WT and OE-*SlBES1*.*8* lines with *P* < 0.01 and *P* < 0.001 respectively (two-tailed Student’s *t*-test).

The decreased leaf complexity in OE-*SlBES1*.*8* plant is reminiscent of the functions of GA in leaf development that increased GA levels or responses could predominantly simplify leaf shape (Jasinski *et al*., 2008; Yanai *et al*., 2011; Lombardi-Crestana *et al*., 2012; Livne *et al*., 2015; Tomlinson *et al*., 2019). To compare the similarity of leaf phenotypic changes caused by OE-*SlBES1*.*8* and GA application, we treated wild type seedlings with GA_3_ and observed the leaf shape. The results showed that WT leaves possessed several serrations in the margin (Fig. 2A). This serration structure can be firstly observed in L6 (the 4^th^ leaf primordium, P4), while no such structure can be found after applying exogenous GA_3_ (Fig. 2A, B). In OE-*SlBES1*.*8* seedlings, there is also no serration structure taken shape (Fig. 2C). Phenotypic observation in mature leaves further confirmed the consistent roles of GA_3_ and SlBES1.8 in leaf development (Supplementary Fig. S4). These results indicate that OE-*SlBES1*.*8* altered the leaf morphogenesis in a manner that similar to GA application.

**Fig. 2.**
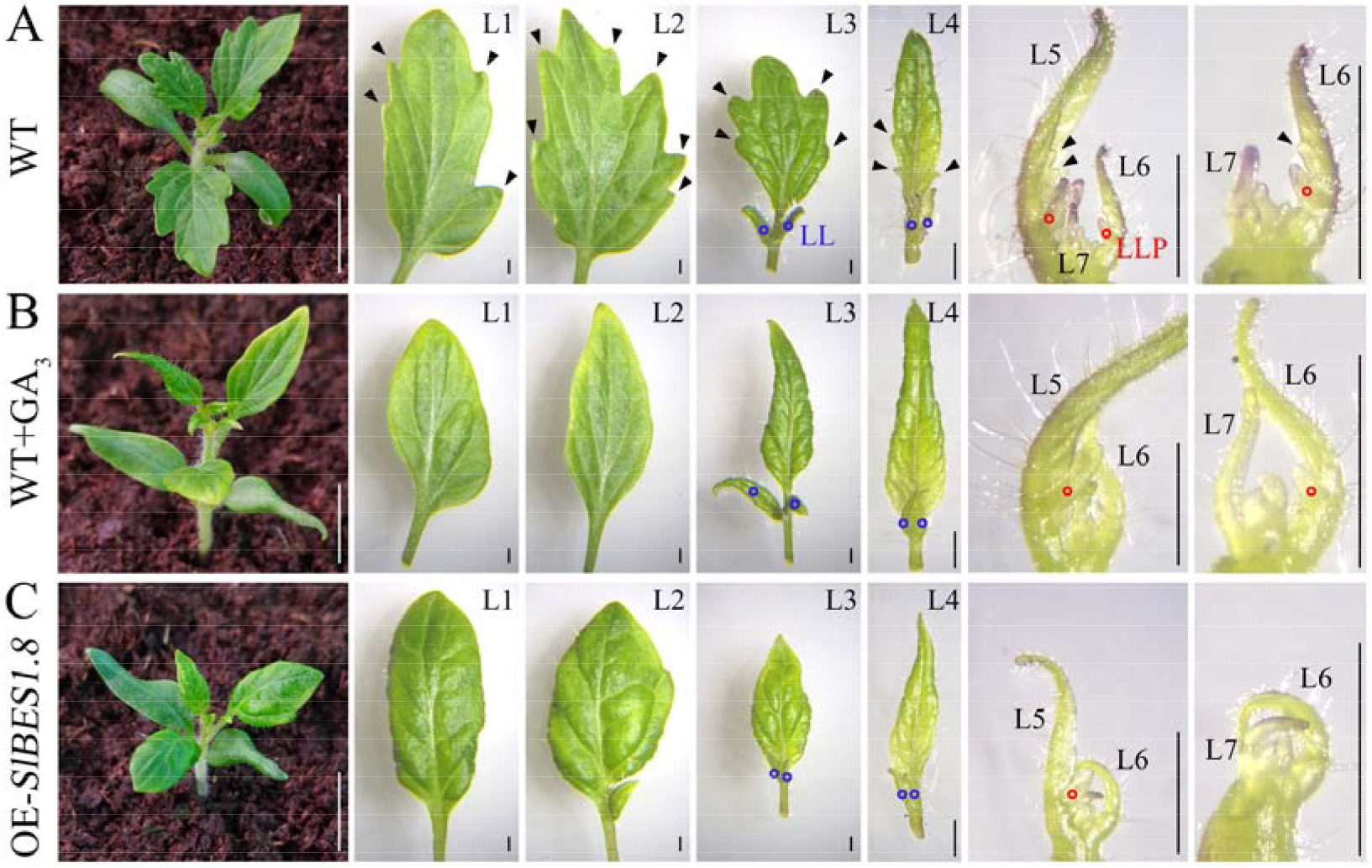
Overexpression of *SlBES1*.*8* resulted in similar consequence as the exogenous application of GA_3_ in leaf morphogenesis. (A) Leaf phenotype of WT seedlings at 16 DPG without GA_3_ application. (B) Leaf phenotype of WT seedlings at 16 DPG with 50 μM GA_3_ application. (C) Leaf phenotype of OE-*SlBES1*.*8* seedlings at 16 DPG without GA_3_ application. Arrowhead indicates the serration of leaf. Blue and red circles indicate lateral leaflet (LL) and lateral leaflet primordium (LLP) respectively. L, leaf; white bar = 1cm; black bar = 1 mm.

### SlBES1.8 decreases the sensibility of tomato to exogenous GA_3_ while increases the sensibility to PAC

The similarity in shaping leaf promotes the possibility that SlBES1.8 may have direct or indirect correlation with GA. We first analyzed the potential *cis*-elements in the promoter of *SlBES1*.*8* and found that there was a GA responsive TATC-box (Fig. 3A). Moreover, qRT-PCR detection verified the induction of GA_3_ to the transcription of *SlBES1*.*8* (Fig. 3B). The primary root of OE-*SlBES1*.*8* seedlings was shorter than WT and was not influenced by GA_3_ treatment (Fig. 3C, D). In addition, the most obvious phenotypic change of tomato seedling after GA_3_ treatment is the elongation of first shoot (Fig. 3C). Surprisingly, the GA-induced first shoot elongation was suppressed in OE-*SlBES1*.*8* lines (Fig. 3C, E). These results suggested that SlBES1.8 decreased the sensibility of tomato seedling to exogenous GA_3_. On the other hand, paclobutrazol (PAC, an inhibitor of GA biosynthesis) treatment could suppress plant growth and produce smaller leaves (Fig. 3F). While the growth suppression of PAC to OE-*SlBES1*.*8* plants was much stronger than WT plants (Fig. 3F). These results indicated that SlBES1.8 increased the sensibility of tomato to PAC. Noticeably, PAC treatment partly recovered the serration in OE-*SlBES1*.*8* leaf margin (Fig. 3F), which further tied SlBES1.8 functions with GA level or response in leaf development.

**Fig. 3.**
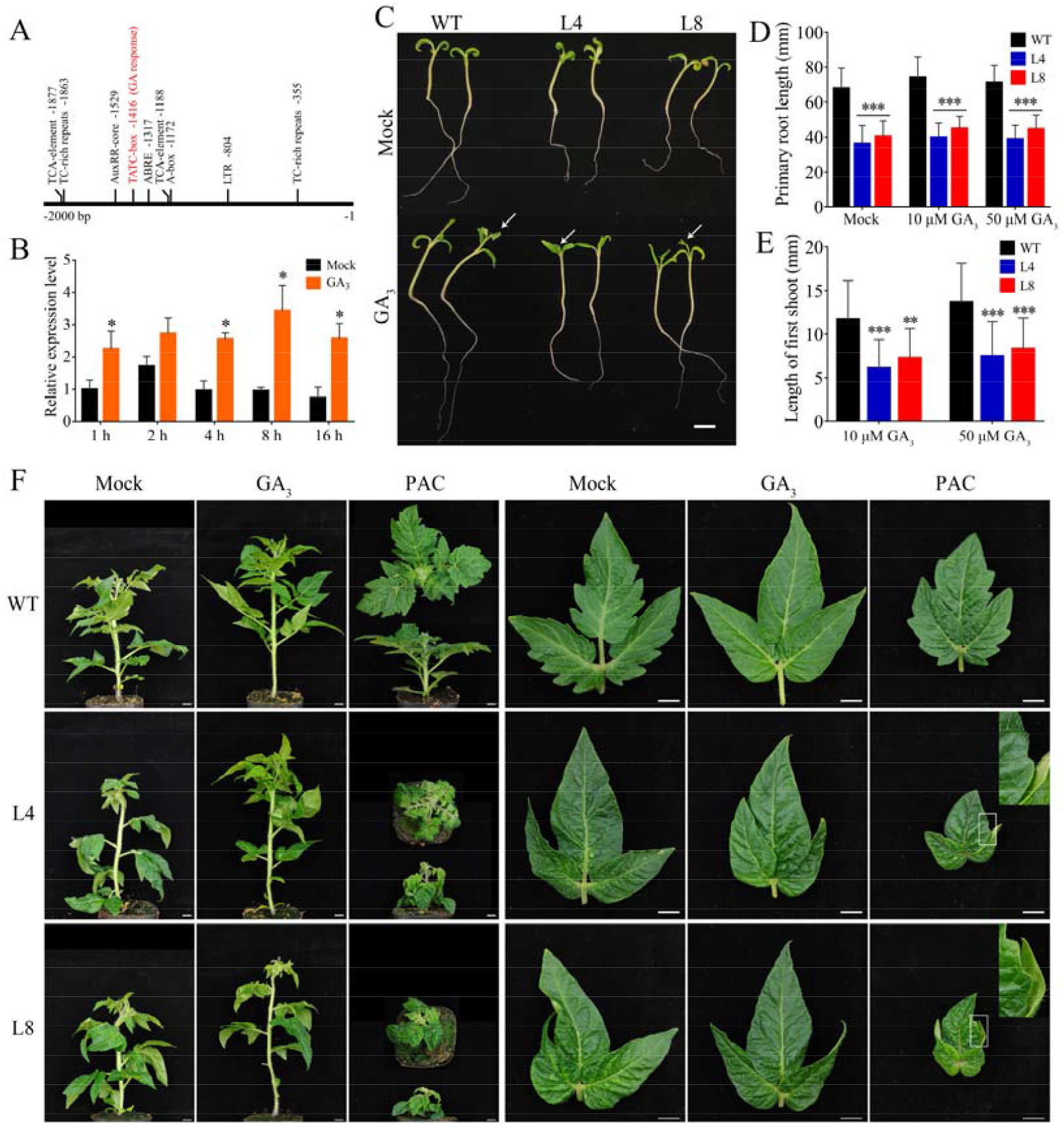
Overexpression of *SlBES1*.*8* reduced the sensitivity to GA_3_ while increased the sensitivity to paclobutrazol (PAC). (A) Potential *cis*-elements in 2 kb upstream promoter of *SlBES1*.*8*. The GA responsive element TATC-box is highlighted with red font. (B) The expression pattern of *SlBES1*.*8* in response to GA_3_ application. Seedlings were soaked into MS/2 medium containing 50 μM GA_3_ for 1, 2, 4, 8 and 16 hours. Values are means ± SD of three biological replicates. (C) Phenotype of WT and OE-*SlBES1*.*8* seedlings at 10 DPG with or without 50 μM GA_3_ application. Arrow indicates the elongated shoot. Bar = 1 cm. (D, E) Length of primary root (D) and first shoot (E) of seedlings with or without GA_3_ application (10 μM and 50 μM). At least 20 seedlings were counted for each column. (F) Phenotype of wild type and OE-*SlBES1*.*8* plants and corresponding leaves with or without GA_3_ (100 μM) or PAC (50 μM) application. Bar = 1 cm. *, ** and *** refer to significant differences with *P* < 0.05, *P* < 0.01 and *P* < 0.001 respectively (two-tailed Student’s *t*-test).

### SlBES1.8 increases endogenous bioactive GA contents in tomato leaf

To assess the effect of OE-*SlBES1*.*8* to GA levels, we detected the endogenous GA contents of WT and OE-*SlBES1*.*8* leaves. According to the GA biosynthesis pathway, we measured the endogenous contents of 16 kinds of GAs in mature leaves, including GA_15_, GA_9_, GA_51_, GA_4_, GA_34_, GA_20_, GA_1_, GA_8_, GA_5_, GA_3_, GA_7_, GA_24_, GA_53_, GA_19_, GA_29_ and GA_6_ (Fig. 4A, B, Supplementary Table S2). Among them, the contents of GA_24_, GA_53_, GA_19_, GA_29_ and GA_6_ were too low to detect. In general, total GAs contents were not significantly changed in OE-*SlBES1*.*8* leaves (Fig. 4C), while the dominant two kinds of GAs, bioactive GA_3_ and inactive GA_8_, were increased and decreased respectively (Fig. 4B, D). Furthermore, the other two bioactive GAs, GA_1_ and GA_4_ were unchanged and increased respectively in OE-*SlBES1*.*8* leaves (Fig. 4E). These results suggested that the increased bioactive GA levels may account for the decreased complexity of OE-*SlBES1*.*8* leaf.

**Fig. 4.**
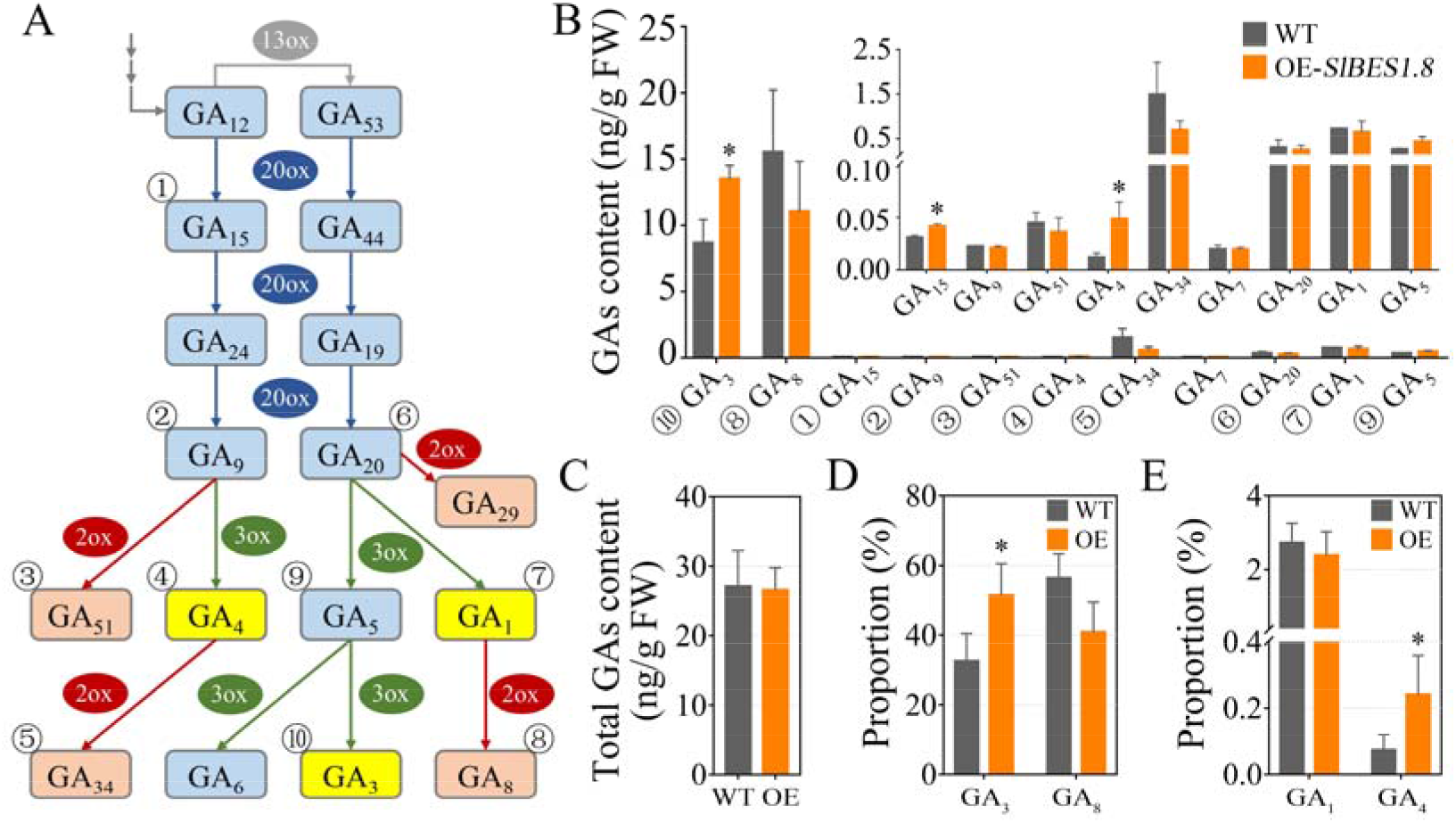
Endogenous gibberellin (GA) contents in WT and OE-*SlBES1*.*8* leaves. (A) Part of generalized scheme of GA biosynthesis and deactivation in higher plants. Serial numbers indicate those GAs that have been detected in our study. 2ox, GA 2-oxidase; 3ox, GA 3-oxidase; 13ox, GA 13-oxidase; 20ox, GA 20-oxidase. (B) Contents of bioactive and inactive GAs detected in our study. (C) Total GAs contents in WT and OE-*SlBES1*.*8* leaves. (D) Proportion of two richest GAs (GA_3_ and GA_8_) in WT and OE-*SlBES1*.*8* leaves. (E) Proportion of another two bioactive GAs (GA_1_ and GA_4_) in WT and OE-*SlBES1*.*8* leaves. Values are means ± SD of three biological replicates. * refers to significant differences between WT and OE-*SlBES1*.*8* line with *P* < 0.05 (two-tailed Student’s *t*-test).

### SlBES1.8 and GA regulate overlapping genomic targets

To view insight of the genome-wide transcriptomic profiling of OE-*SlBES1*.*8* plants, we performed RNA-sequencing for WT and OE-*SlBES1*.*8* leaf primordium at P4 stage. With the criterions of fold change (FC) > 2 and adjust *P*-value (*Q*) < 0.05, a total of 259 and 297 differentially expressed genes (DEGs) were influenced by GA_3_ application and OE-*SlBES1*.*8* respectively (Fig. 5A, Supplementary Table S3, 4). Among them, 42 DEGs were regulated by both GA_3_ and SlBES1.8, which promoted the close relationship between GA_3_ and SlBES1.8 (Fig. 5B, Supplementary Table S5). If SlBES1.8 mimics the functions of GA in leaf development, SlBES1.8 should affect the expression of genes that influenced by GA in a similar manner. Indeed, by analyzing the expression profiles of these 42 co-DEGs, we found that 36 DEGs (85.7%) were affected in the same way by GA_3_ and OE-*SlBES1*.*8* (Fig. 5C, group □), whereas only six DEGs were influenced in the opposite way (Fig. 5C, group □). Notably, when compared the expression of these co-DEGs in GA_3_ treated OE-*SlBES1*.*8* leaves with mocked WT leaves (OE GA_3_ vs WT Mock), the expression changes displayed a superimposed effect of OE-*SlBES1*.*8* and GA_3_ treatment, which suggested that on the basis of exogenous GA_3_ application, OE-*SlBES1*.*8* further strengthened GA signaling (Fig. 5C). Meanwhile, by comparing the expression of these 42 DEGs in WT and OE-*SlBES1*.*8* leaves after GA_3_ treatment, we found that 28 DEGs (66.7%, group □) exhibited weaker expression changes in OE-*SlBES1*.*8* leaves than that in WT (Fig. 5D), suggesting that OE-*SlBES1*.*8* repressed their responses to GA_3_ application, which further supported our previous conclusion that SlBES1.8 decreased the sensibility of tomato to exogenous GA_3_. Collectively, OE-*SlBES1*.*8* and GA_3_ application influenced a mass of common targets in the similar manner.

**Fig. 5.**
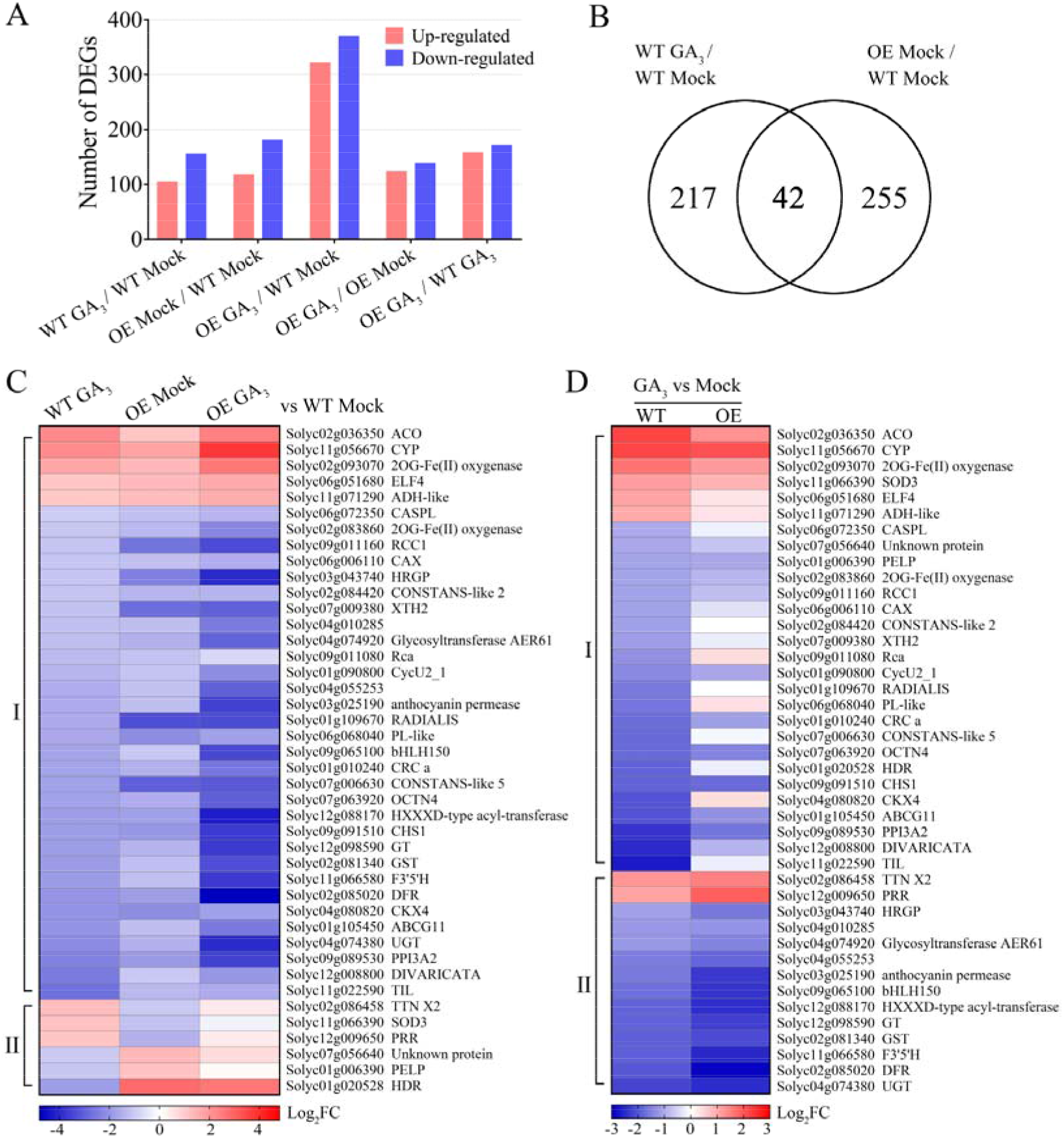
Transcriptome analysis of WT and OE-*SlBES1*.*8* leaves with or without GA_3_ application. (A) Numbers of up-or down-regulated differentially expressed genes (DEGs, FC > 2, *Q* < 0.05) in distinct comparisons. (B) Venn diagram showing the overlap between sets of DEGs influenced by GA_3_ treatment in WT (WT GA_3_ / WT Mock) or by overexpression of *SlBES1*.*8* (OE Mock / WT Mock). (C) Heat map showing the expression profiles of 42 common DEGs in (B). Group □ containing DEGs that are both up-or down-regulated by GA_3_ and OE-*SlBES1*.*8*. Group □ containing DEGs that are regulated by GA_3_ and OE-*SlBES1*.*8* with opposite trends. (D) Expression comparison of those 42 common DEGs after GA_3_ application in WT and OE-*SlBES1*.*8* leaves. Group □ and □ containing DEGs with smaller or bigger effect respectively by GA_3_ application in OE-*SlBES1*.*8* leaves compared with WT. FC, fold change.

### SlBES1.8 binds to and represses the activities of SlGA2ox2, SlGA2ox6 and SlGID1b-1 promoters

To elucidate the regulatory mechanism of SlBES1.8 in regulating leaf development by mediating GA levels or responses, we carefully analyzed all DEGs affected by OE-*SlBES1*.*8* and GA_3_ application, and found that merely none of them was reported to involve in GA biosynthesis or signaling (Supplementary Table S3, 4). This may be caused by the very early developmental status of RNA-seq samples (the 4^th^ leaf primordium of seedlings at 16DPG) that few of genes exhibited differential expression in this stage. Hence, we selected to explore the potential targets of SlBES1.8 by using the older leaves (that is L3 presented in Fig. 2) as the materials.

The increased bioactive GA contents could be caused by the up-regulation of GA biosynthetic genes or the down-regulation of GA deactivated genes. Given that SlBES1.8 acts as a transcriptional repressor (Su *et al*., 2021), we mainly considered the repression of SlBES1.8 to the expression and responsiveness of *GA 2-oxidase* (*GA2ox*) genes, which are the best-characterized GA deactivated genes. The relative expressions of nine tomato *GA2ox* genes in WT and OE-*SlBES1*.*8* leaves with or without GA_3_ treatment were explored by qRT-PCR. Overall, *SlGA2oxs* transcriptions were greatly induced by GA_3_ in WT leaves as expected (Fig. 6A, B, Supplementary Fig. S5A). Among them, the expression levels of *SlGA2ox2* and *SlGA2ox6* were significantly reduced in OE-*SlBES1*.*8* leaves, and their responses to GA_3_ application were also suppressed by OE-*SlBES1*.*8* (Fig. 6A, B), suggesting that SlBES1.8 may increase the bioactive GA contents through repressing the expressions of *SlGA2ox2* and *SlGA2ox6*. Meanwhile, we also detected the relative expression of genes involved in GA signaling (Fig. 6C, Supplementary Fig. S5B). Among them, *SlGID1b-1* showed reduced expression in OE-*SlBES1*.*8* leaves (Fig. 6C), indicating a potential direct regulation by SlBES1.8.

**Fig. 6.**
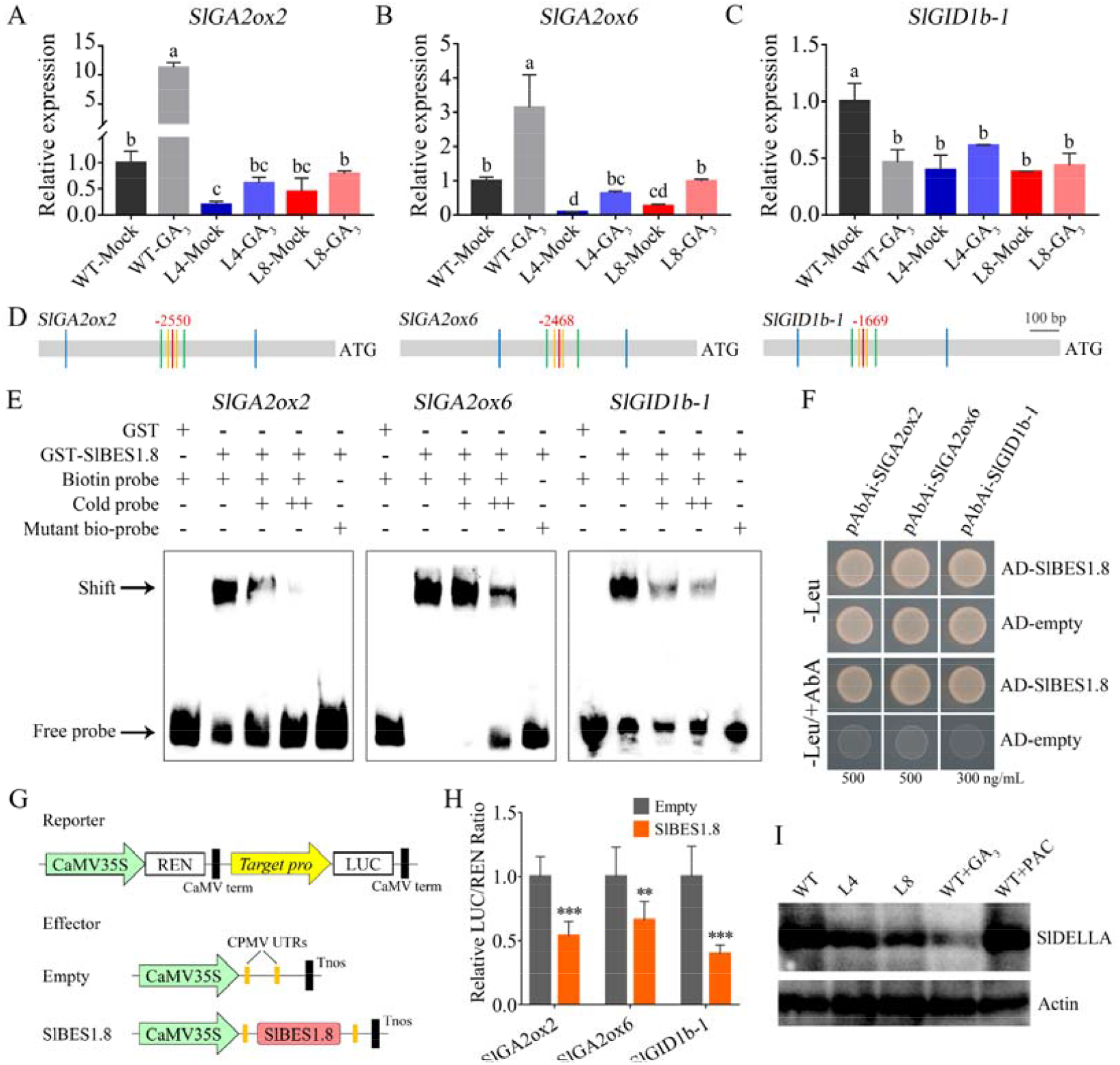
SlBES1.8 directly represses the expression of *SlGA2ox2, SlGA2ox6* and *SlGID1b-1* and promotes the degradation of SlDELLA. (A-C) Relative expression levels of *SlGA2ox2* (A), *SlGA2ox6* (B) and *SlGID1b-1* (C) in response to GA3 application in WT and OE-*SlBES1*.*8* lines. Values are means ± SD of three biological replicates. Different letters indicate significant differences (*P* < 0.05) according to one-way ANOVA test. (D) Diagram showing the promoter fragments of *SlGA2ox2, SlGA2ox6* and *SlGID1b-1* used in electrophoretic mobility shift assay (EMSA, between orange lines), yeast one-hybrid assays (Y1H, between green lines) and dual-luciferase assays (between blue lines). The position of G-box (CACGTG) is indicated by red line. (E) EMSA showing the binding of SlBES1.8 to the promoters of *SlGA2ox2, SlGA2ox6* and *SlGID1b-1*. GST alone incubated with biotin probe was used as negative control. Mutant biotin probe and competitor probe confirmed the specific binding. ++, increased amount of competitor probes. (F) Y1H showing the binding of SlBES1.8 to the promoters of *SlGA2ox2, SlGA2ox6* and *SlGID1b-1*. Yeast transformants were cultured on SD/-Leu or SD/-Leu/+AbA mediums for 3 days. pGADT7-empty plasmid was used as control. The screening concentration of AbA was indicated in the bottom. (G) Diagram of reporters and effector used in dual-luciferase assays. Promoters of *SlGA2ox2, SlGA2ox6* and *SlGID1b-1* were constructed into reporter, and full length coding sequence of *SlBES1*.*8* was constructed into effector. (H) Results of dual-luciferase assays showing the inhibition of SlBES1.8 to the promoter activity of *SlGA2ox2, SlGA2ox6* and *SlGID1b-1*. The LUC/REN ratio of the group of empty effector was regarded as calibrator (set as 1). At least six biological replicates were used for each column. ** and *** refer to significant differences between empty and SlBES1.8 effectors with *P* < 0.01 and *P* < 0.001 respectively (two-tailed Student’s *t*-test). (I) Immunoblots showed the content of SlDELLA. GA_3_ promoted the degradation whereas PAC promoted the accumulation of SlDELLA, as expected. Actin is used as a loading control.

To confirm the direct regulation of SlBES1.8 to *SlGA2ox2, SlGA2ox6* and *SlGID1b-1*, we conducted electrophoretic mobility shift assay (EMSA), yeast one-hybrid assay (Y1H) and dual-luciferase assay according to the existence of G-box (CACGTG), a well-identified binding motif of BES1 family (He *et al*., 2005; Yin *et al*., 2005), in the promoters of *SlGA2ox2, SlGA2ox6* and *SlGID1b-1* (Fig. 6D). EMSA results showed the specific binding of SlBES1.8 to the selected promoter fragments (Fig. 6E). Y1H results further confirmed the binding of SlBES1.8 to *SlGA2ox2, SlGA2ox6* and *SlGID1b-1* promoters (Fig. 6F). Moreover, we used dual-luciferase system to detect the effects of SlBES1.8 to the promoter activities in tobacco leaves and found that SlBES1.8 significantly repressed the activities of *SlGA2ox2, SlGA2ox6* and *SlGID1b-1* promoters (Fig. 6G, H). The transcriptional repression of SlBES1.8 to *SlGA2ox2* and *SlGA2ox6* may be the reason of increased bioactive GA contents in OE-*SlBES1*.*8* leaves, which should promote the degradation of SlDELLA protein. Indeed, by performing immunoblot, we confirmed the decreased level of SlDELLA protein in OE-*SlBES1*.*8* lines (Fig. 6I). Collectively, these results indicated that SlBES1.8 increased the endogenous bioactive GA levels by repressing the expression of *SlGA2ox2* and *SlGA2ox6*, which further promoted the degradation of SlDELLA protein, resulting in a similar effect with increased GA levels or responses in tomato leaf development.

### SlDELLA physically interacts with SlBES1.8 and inhibits its transcriptional regulation ability by abolishing SlBES1.8-DNA binding

DELLA protein lacks a DNA-binding domain, which promotes the necessary of intermediate proteins for its functions (Yoshida *et al*., 2014; Marín-de la Rosa *et al*., 2015). In Arabidopsis, DELLAs were proved to physically interact with BES1/BZR1, which further inhibited the DNA binding ability of BZR1 (Bai *et al*., 2012; Gallego-Bartolomé *et al*., 2012; Li *et al*., 2012). The characteristic domains of DELLA proteins were conserved in tomato and Arabidopsis (Supplementary Fig. S6), which raised the possibility that tomato DELLA protein may interact with SlBES1 transcription factors in a similar manner, including SlBES1.8. To prove this assumption, we investigated the protein-protein interaction between SlDELLA and SlBES1.8 by yeast two-hybrid assay (Y2H), bimolecular fluorescence complementation (BiFC) assay and firefly luciferase complementation imaging (LCI) assay. The results showed that SlDELLA can truly interact with SlBES1.8 (Fig. 7A-C). Furthermore, we explored the interacted domains of SlDELLA and SlBES1.8 by generating a series of deletion constructs used in LCI assays according to the amino acid sequence analysis (Supplementary Fig. S7A). The results showed that the deletion of PEST domain of SlBES1.8 was sufficient to prevent the interaction with SlDELLA (Supplementary Fig. S7B), and deletion of DELLA motif of SlDELLA abolished the interaction with SlBES1.8 (Supplementary Fig. S7C), indicating that the PEST domain of SlBES1.8 and the DELLA motif of SlDELLA were critical for their protein-protein interaction.

**Fig. 7.**
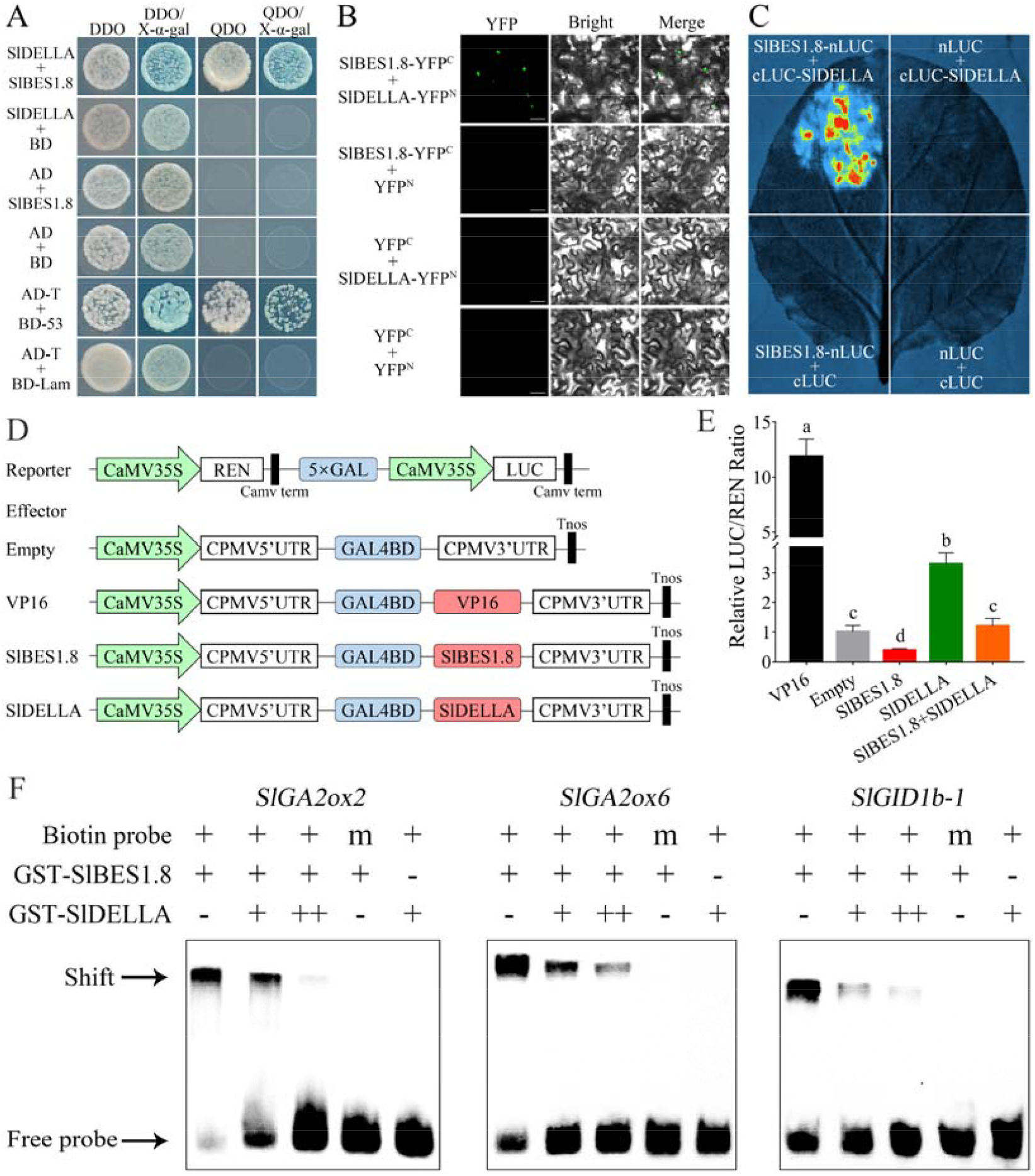
SlDELLA interacts with SlBES1.8 and suppresses its transcriptional repression ability by inhibiting SlBES1.8-DNA binding. (A) Yeast two-hybrid (Y2H) assay for interaction between SlDELLA and SlBES1.8 proteins. AD-T plus BD-53 or BD-Lam were used as the positive and negative control respectively. DDO, SD/-Leu/-Trp medium; QDO, SD/-Ade/-His/-Leu/-Trp medium. (B) Bimolecular fluorescence complementation (BiFC) assay in tobacco leaves for interaction between SlBES1.8 and SlDELLA proteins. Bar = 50 μm. (C) Firefly luciferase complementation imaging (LCI) assay in tobacco leaves for interaction between SlBES1.8 and SlDELLA proteins. (D) Diagram of reporter and effectors used in dual-luciferase assays. *SlDELLA* and *SlBES1*.*8* were fused with GAL4BD to generate the effector constructs. VP16 was used as the transcriptional activation control. (E) Suppression of SlDELLA on the transcriptional repression activity of SlBES1.8 detecting by GAL4-responsive reporter system in tobacco leaves. Reporter was co-expressed with SlBES1.8 or SlDELLA effector alone or together in tobacco leaves. The LUC/REN ratio of the group of empty effector was regarded as calibrator (set as 1). At least six biological replicates were used for each column. The significant differences are indicated with letters (*P* < 0.05, one-way ANOVA test). (F) EMSA showing the inhibition of SlDELLA on the binding of SlBES1.8 to the promoters of *SlGA2ox2, SlGA2ox6* and *SlGID1b-1*. Biotin probes were incubated with GST-SlBES1.8 alone or together with GST-SlDELLA protein. ++ indicates increased amount of GST-SlDELLA proteins; m, mutant biotin probe.

We next analyzed the effect of SlDELLA interaction to the transcriptional regulation activity of SlBES1.8 by GAL4-responsive reporter system in tobacco leaves. Full length of *SlDELLA* and *SlBES1*.*8* coding sequences were fused with GAL4BD respectively to generate the effectors, which were subsequently co-expressed with reporter in tobacco leaves (Fig. 7D). As the transcriptional activation control, VP16 greatly induced the expression of *LUC*, indicating the effectiveness of this system. In line with our previous detection (Su *et al*., 2021), SlBES1.8 significantly repressed *LUC* expression, suggesting the transcriptional repression activity. On the contrary, SlDELLA increased *LUC* expression and exhibited transactivation activity, which was consistent with the rice DELLA protein, SLR1 (Hirano *et al*., 2012). Dramatically, SlDELLA antagonized the transcriptional regulation activity of SlBES1.8 when they co-expressed together (Fig. 7E). Besides, using EMSA, we confirmed that the bindings of SlBES1.8 to the promoter fragments of *SlGA2ox2, SlGA2ox6* and *SlGID1b-1* were suppressed by the presence of SlDELLA (Fig. 7F). Noticeably, SlDELLA was unable to bind these fragments in the absence of SlBES1.8, as expected. Taken together, these results indicated that through protein-protein interaction, SlDELLA can inhibit the transcriptional regulation activity of SlBES1.8 by repressing its DNA binding ability.

## Discussion

Leaf morphological diversity is achieved by fine-tuning the morphogenetic window during development that prolonged window resulted in complex leaf patterning whereas shortened window led to simpler leaf shape (Kierzkowski *et al*., 2019). During this elaborated developmental process, several regulators acted coordinately to affect different aspects of leaf shape, ensuring the establishment of a suitable pattern (Du *et al*., 2018; Israeli *et al*., 2021). Hormones play critical roles in various aspects of plant development including leaf formation (Shwartz *et al*., 2016). Among them, GA is known to reduce the complexity of leaf pattern in tomato. Tomato plants treated with exogenous GA or mutated in *DELLA* locus exhibited simplified leaf that with less leaflet production and smooth margin (Jasinski *et al*., 2008; Lombardi-Crestana *et al*., 2012; Livne *et al*., 2015; Shwartz *et al*., 2016; Tomlinson *et al*., 2019). By investigating the early leaf development of tomato DELLA mutant *pro*, the simplification of *pro* leaf was caused by the combination of accelerated early developmental growth and delayed leaflet initiation (Jasinski *et al*., 2008), indicating the effects of increased GA signaling in shortening leaf morphogenetic phase. Since the clarification of GA functions in leaf development, several regulators that can greatly influence leaf morphogenesis were reported to fulfill their roles by affecting GA homeostasis, such as LA and KNOXI (Jasinski *et al*., 2008; Yanai *et al*., 2011). In this research, we also identified a tomato leaf development regulator, SlBES1.8, that promoted leaf simplification by mediating GA homeostasis and signaling. We offered multiple lines of evidence to support this finding. First, OE-*SlBES1*.*8* lines in micro-tom and *Ailsa Craig* background presented less leaflet with smooth margin, which was highly resemble to GA-treated wild type leaves. Second, endogenous bioactive GA levels were increased in OE-*SlBES1*.*8* leaves, which subsequently led to the degradation of SlDELLA as expected. Third, OE-*SlBES1*.*8* and GA affected a set of overlapping targets in a similar manner. Fourth, SlBES1.8 directly regulated the expression of *SlGA2ox2, SlGA2ox6* and *SlGID1b-1*. And fifth, SlBES1.8 physically interacted with SlDELLA and were inhibited by such interaction in DNA binding ability. In summary, we proposed a working model to depict the molecular mechanism of SlBES1.8 in regulating tomato leaf morphogenesis by mediating GA deactivation and signaling (Fig. 8). Although we determined the correlation of SlBES1.8 with GA, the downstream targets of GA signaling that directly contribute to leaf development remain largely unknown and need to be investigated. Also, new modifiers of leaf pattern that affect variable aspects of leaf shape such as lobe/serration, leaflet number and size, are worth to be discovered.

**Fig. 8.**
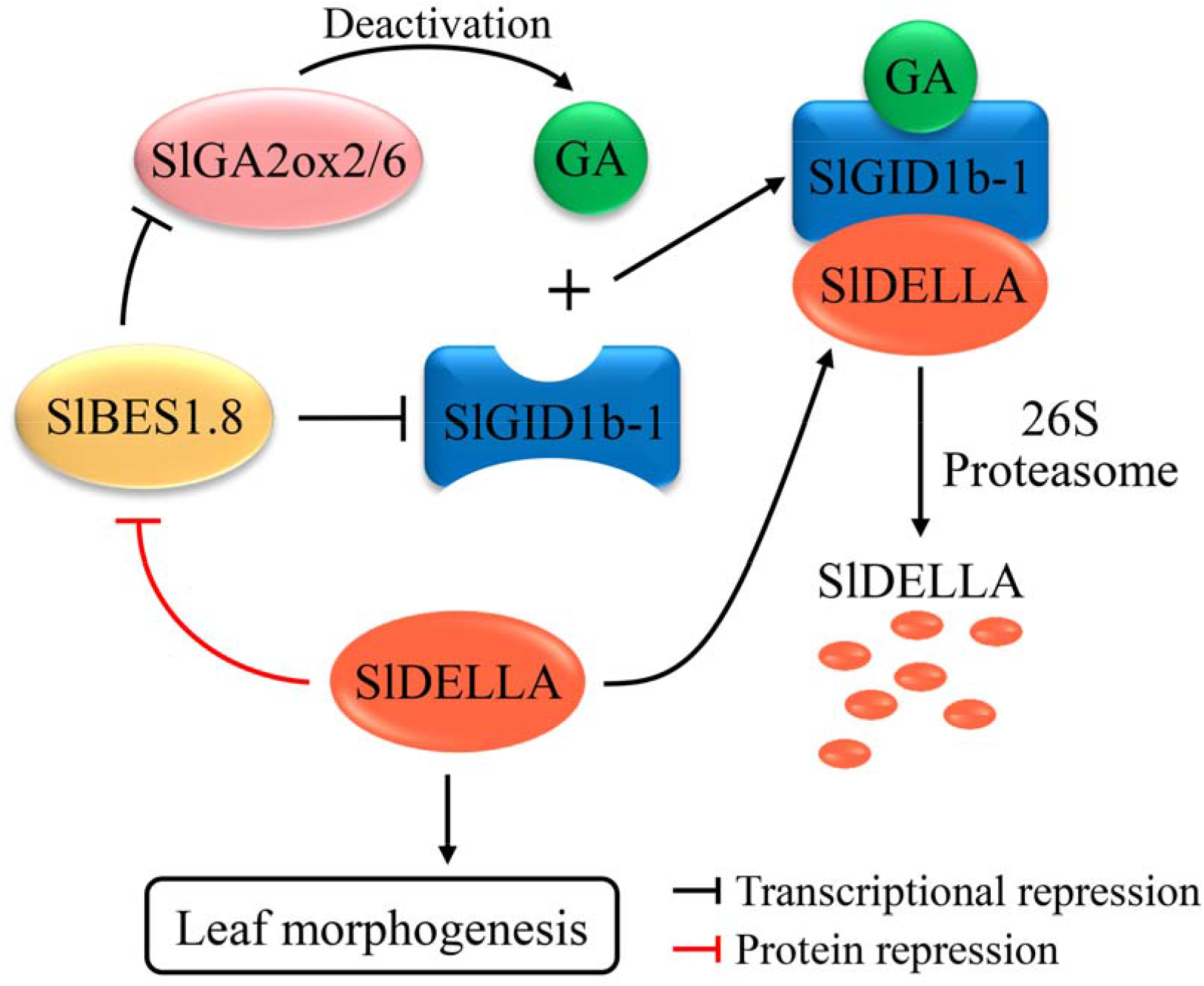
Proposed model depicting the molecular mechanism of SlBES1.8 in the regulation of tomato leaf morphogenesis. SlBES1.8 represses the transcriptions of *SlGA2ox2* and *SlGA2ox6*, which encode GA deactivation enzymes, to positively regulate bioactive GA contents and SlDELLA degradation. Meanwhile, SlBES1.8 also represses the expression of GA receptor, *SlGID1b-1*, leading to a feedback regulation in SlDELLA degradation. On the other hand, SlDELLA physically interacts with SlBES1.8 and inhibits its transcriptional regulation ability. The degradation of SlDELLA releases its inhibition to SlBES1.8, which further promotes SlDELLA degradation. By such mediation both in GA level and signaling, SlBES1.8 greatly influences tomato leaf morphogenesis.

To identify the functions of SlBES1.8, we also generated CRISPR/Cas9 mutants for it and expected the corresponding phenotypic changes to OE-*SlBES1*.*8* plants in leaf development. We selected two homozygous lines with distinct mutations to observe the leaf phenotype and failed to obtain obvious developmental differences (Supplementary Fig. S3). Two reasons may explain this. First, the constitutive expression level of *SlBES1*.*8* is very low, with an average of 0.86 TPM transcription in wild type leaves, as a contrast, its transcription was increased to 1555.67 TPM in OE-*SlBES1*.*8* leaves (Supplementary Table S4). The low content of SlBES1.8 contributed to a weak repression to *SlGA2ox2* and *SlGA2ox6* transcription that its knockout was not enough to affect endogenous GA levels. Second, decreasing GA levels or responses contributed a relative minor effect on leaf complexity. Tomato mutants with reduced GA levels or responses generally sustained pleiotropic effects, which made it difficult to interpret its effect on leaf formation (Koornneef *et al*., 1990; Jasinski *et al*., 2008; Yanai *et al*., 2011; Livne *et al*., 2015; Tomlinson *et al*., 2019). Besides, specific reducing endogenous GA levels in leaf by overexpressing GA deactivated gene *GA2ox* under the driving of leaf-specific promoter *pFIL* resulted in more complex leaf pattern, while was moderate compared to the substantial simplification effect caused by increasing GA levels or responses (Shwartz *et al*., 2016).

BR and GA are known as two principal groups of plant growth-promoting phytohormones for their functions in promoting cell elongation and differentiation. Defects in either of their biosynthesis or signaling transduction could result in inhibited plant growth and dwarfism (Schwechheimer, 2011; Fridman and Savaldi-Goldstein, 2013). The overlapping functions of BR and GA implied the potential crosstalk between them, which attracted extensive attentions in the recent one decade (Li and He, 2013). A breakthrough was the signaling interaction between these two hormones reported by three different laboratories at the same time. In this signaling model, the negative regulator of GA signaling, DELLAs, can physically interact with BES1 and BZR1, the positive regulators of BR signaling (Bai *et al*., 2012; Gallego-Bartolomé *et al*., 2012; Li *et al*., 2012). Specifically, DELLAs only interacted with the dephosphorylated active BZR1 and further inhibited BZR1-DNA binding ability (Bai *et al*., 2012). This DELLA-BES1/BZR1 interaction also existed in our research that tomato DELLA protein can physically bind to SlBES1.8 to interfere with its function by abolishing SlBES1.8-DNA binding (Fig. 7). Later, the synthesis model was proposed based on the results obtained from Arabidopsis and rice that BR regulated the biosynthesis of GA. In Arabidopsis, the expressions of multiple GA biosynthesis genes *GA20oxs* and *GA3oxs* were impaired in BR mutants. Correspondingly, the bioactive GA contents were reduced, which was partly responsible for the growth defects of BR mutants because external GA application or expression of *GA20ox1* partially restored the defects. Furthermore, the direct regulation of BES1 to the promoters of *GA20ox1, GA3ox1* and *GA3ox4* were verified (Unterholzner *et al*., 2015). Similarly, BZR1 directly bound to the promoters of *GA20ox2, GA3ox2* and *GA2ox3* to regulate their expression in rice (Tong *et al*., 2014). In our study, we determined the direct repression of SlBES1.8 to *SlGA2ox2* and *SlGA2ox6*, which led to the increase of bioactive GA contents that resulted in a similar phenotype to increased GA level or response. Taken together, our results obtained from tomato, in a certain extent, expanded our knowledge to BR and GA crosstalk both in signaling level and biosynthesis level, and more efforts need to be made in elaborating the crosstalk of these two hormones in different species.

As the key regulators of BR signaling, *BES1* genes involved in numerous biological processes by directly interacting with key elements from other pathways. The functional identification of *BES1* family genes were widely investigated in Arabidopsis and other species, with critical roles in regulating cell elongation and division (Xie *et al*., 2011), stress tolerance (Nolan *et al*., 2017), immune signaling (Kang *et al*., 2015), meristem differentiation (Kondo *et al*., 2014) and male sterility (Chen *et al*., 2019). Particularly, in tomato, the functional investigations of this family were mainly focused on the two key members, SlBES1 and SlBZR1. With convincing evidences, SlBES1 and SlBZR1 were verified to play important roles in regulating heat stress tolerance (Yin *et al*., 2018), mediating autophagosome formation (Wang *et al*., 2019), affecting pollen development (Yan *et al*., 2020), influencing wood formation (Lee *et al*., 2021), releasing apical dominance (Xia *et al*., 2021) and promoting fruit softening (Liu *et al*., 2021), while little is known about the functions of other members. Our findings that SlBES1.8 greatly influences tomato leaf morphogenesis obviously enriched the functional diversity of tomato *BES1* family, and more functional analyses of this family members in tomato and other species would be benefit to understand the crosstalk of BR with other pathways in regulating plant growth and stress responses.

## Supplementary data

Supplementary data are available at *JXB* online.

***Fig. S1***. Relative mRNA level of *SlBES1*.*8* in over-expressed lines.

***Fig. S2***. Phenotype of one-month-old WT and OE-*SlBES1*.*8* plants in *Ailsa Craig* background.

***Fig. S3***. Knockout of *SlBES1*.*8* didn’t influence the normal development of tomato leaf.

***Fig. S4***. Overexpression of *SlBES1*.*8* resulted in similar consequence as the exogenous application of GA_3_ in leaf morphogenesis.

***Fig. S5***. Expression pattern of genes involved in GA deactivation and signaling in WT and OE-*SlBES1*.*8* leaves in response to GA_3_ application.

***Fig. S6***. Amino acids alignment of tomato and Arabidopsis DELLA proteins by ClustalX.

***Fig. S7***. The PEST domain of SlBES1.8 and DELLA motif of SlDELLA are critical for their interaction.

***Table S1***. Primers used in this study.

***Table S2***. GAs contents in WT and OE-*SlBES1*.*8* leaves.

***Table S3***. DEGs after GA_3_ treatment in WT leaves.

***Table S4***. DEGs resulted by OE-*SlBES1*.*8*.

***Table S5***. 42 DEGs influenced by both GA_3_ and OE-*SlBES1*.*8*.

## Acknowledgements

We would like to thank Analytical and Testing Center of Chongqing University for providing laser scanning confocal microscope analysis. This work was supported by the Fundamental Research Funds for the Central Universities (No. 2021CDJZYJH-002) and the National Natural Science Foundation of China (No. 31972470, 31772370, 32002100).

## Author contributions

Z. L. supervised the research; Z. L., Z. X. and D. S. designed the experiments; D. S., W. X., Q. L. and L. W. performed the experiments; Y. L. provided experimental assistance; D. S. wrote the manuscript and Z. X., Y. L. and Y. S. revised the paper. All authors had read and approved the final manuscript.

## Data availability

All data supporting the results of this study are included in the paper and its supplementary files.

